# Action-based predictions affect visual perception, neural processing, and pupil size, regardless of temporal predictability

**DOI:** 10.1101/2021.02.11.430717

**Authors:** Christina Lubinus, Wolfgang Einhäuser, Florian Schiller, Tilo Kircher, Benjamin Straube, Bianca M. van Kemenade

## Abstract

Sensory consequences of one’s own action are often perceived as less intense, and lead to reduced neural responses, compared to externally generated stimuli. Presumably, such sensory attenuation is due to predictive mechanisms based on the motor command (efference copy). However, sensory attenuation has also been observed outside the context of voluntary action, namely when stimuli are temporally predictable. Here, we aimed at disentangling the effects of motor and temporal predictability-based mechanisms on the attenuation of sensory action consequences. During fMRI data acquisition, participants (N = 25) judged which of two visual stimuli was brighter. In predictable blocks, the stimuli appeared temporally aligned with their button press (active) or aligned with an automatically generated cue (passive). In unpredictable blocks, stimuli were presented with a variable delay after button press/cue, respectively. Eye tracking was performed to investigate pupil-size changes and to ensure proper fixation. Self-generated stimuli were perceived as darker and led to less neural activation in visual areas than their passive counterparts, indicating sensory attenuation for self-generated stimuli independent of temporal predictability. Pupil size was larger during self-generated stimuli, which correlated negatively with blood oxygenation level dependent (BOLD) response: the larger the pupil, the smaller the BOLD amplitude in visual areas. Our results suggest that sensory attenuation in visual cortex is driven by action-based predictive mechanisms rather than by temporal predictability. This effect may be related to changes in pupil diameter. Altogether, these results emphasize the role of the efference copy in the processing of sensory action consequences.

## 1. Introduction

The sensory consequences of one’s own actions result in a less intense experience than identical but externally generated events. This phenomenon, called sensory attenuation (Brown et al., 2013; Hughes et al., 2013), is thought of as one of the mechanisms allowing an organism to distinguish between internal and external events (Frith et al., 2000; Haggard and Tsakiris, 2009). Attenuation of self-generated sensory events is a phenomenological experience (Blakemore et al., 1999a; Sato, 2008; Weiss et al., 2011) reflected in reduced neural processing (Schafer and Marcus, 1973; Blakemore et al., 1998, 1999b; Martikainen, 2004; Bäß et al., 2008; Aliu et al., 2009; Shergill et al., 2013; Straube et al., 2017; Arikan et al., 2019; Pazen et al., 2019; Uhlmann et al., 2020; Schmitter et al., 2021). These findings have predominantly been discussed within the framework of internal forward models which suggests that the sensory consequences of one’s actions are predicted based on efference copies generated during motor planning (Wolpert et al., 1995). Despite the efference-copy hypothesis being suggested as a general mechanism occurring in all modalities (Brown et al., 2013), sensory attenuation has mainly been investigated in somatosensation (Blakemore et al., 1998, 1999a, 1999b) and audition (Sato, 2008; Aliu et al., 2009; Bäß et al., 2009; Weiss et al., 2011; Sanmiguel et al., 2013; Mifsud et al., 2016a), with evidence in the visual domain remaining inconclusive. While some studies reported lower perceptual thresholds (Cardoso-Leite et al., 2010; Dewey and Carr, 2013) and/or reduced neural responses (Leube, 2003; Straube et al., 2017; Arikan et al., 2019; Pazen et al., 2019; Uhlmann et al., 2020; Schmitter et al., 2021) for self-generated visual stimuli, others observed no (Schwarz et al., 2018) or ambiguous effects (Yon and Press, 2017).

The notion of motor action being crucial for sensory attenuation is challenged by the observation that mere temporal prediction of a stimulus can also attenuate perceptual and neural processing (Summerfield et al., 2008; Bendixen et al., 2009; Alink et al., 2010; Todorovic et al., 2011; Kok et al., 2012; John-Saaltink et al., 2015). It has been suggested that in addition to motor predictions, better temporal predictability of stimulus onset may contribute to sensory attenuation and neural suppression effects for actively generated stimuli, simply because of a heightened temporal control when the presentation of a stimulus is caused by one’s own action (Hughes et al., 2013). Many paradigms indeed compare self-generated stimuli with stimuli that are externally generated at random time points, which creates a confound as the temporal predictability between actively and externally generated stimuli differs in such paradigms. As such, sensory attenuation for self-generated stimuli may not solely be due to efference-copy mechanisms but be explained at least partially by increased temporal predictability of these stimuli.

This study aimed to disentangle the roles of action-based and general temporal prediction mechanisms in sensory attenuation of visual action consequences. During functional magnetic resonance imaging (fMRI), participants engaged in a visual intensity judgment task, judging which of two subsequent stimuli was brighter. Stimuli were elicited either by an active button press or by the computer; stimulus onset was either temporally predictable or unpredictable. We manipulated luminance, the physical quantity related to the intensity of a visual stimulus, and considered brightness, representing the *perceived* intensity as dependent variable. This was motivated by the extensive work on sensory attenuation in the auditory (e.g., Sato, 2009; Weiss et al., 2011) and somatosensory (Blakemore et al., 1998, 1999a) senses. The results of these lines of research suggest that perceived stimulus intensity is attenuated for sensory events resulting from one’s own actions. Neurons in visual cortex have long been known to be sensitive to orientation (Graham et al., 1993; Ling et al., 2009) and contrast (Boynton et al., 1996; Avidan et al., 2002), and were thought of as less (if at all) responsive to uniform illumination (Hubel and Wiesel, 1968). However, more recent evidence suggests that primary visual areas in humans can respond to homogenous luminance (Penacchio et al., 2013) and changes in luminance (Haynes et al., 2004; Vinke and Ling, 2020) (but also see: Cornelissen, 2006). In the light of this, we expected that visual events would be perceived as darker and result in a suppressed BOLD response in primary visual cortex for active as compared to passive trials. Furthermore, we hypothesized that temporal predictability of stimulus onset and voluntary action would interact such that the attenuation (behavioral and neural) was strongest for trials being active and predictable.

Finally, we were interested in the role of pupil size in sensory attenuation effects. It is long established that pupil size is influenced by stimulus luminance (Loewenfeld, 1958; Larsen and Waters, 2018). However, pupil size is also affected by brightness such that stimuli perceived or expected as brighter give rise to stronger constriction, even if luminance remains physically unchanged (Laeng and Endestad, 2012; Binda et al., 2013; Naber and Nakayama, 2013). Thus, we expected that dilation was predictive of perceived stimulus intensity (i.e., lower perceived intensity resulting in bigger pupil size). In addition, we aimed to explore the relation between pupil size and the neural effects of sensory attenuation.

## 2. Materials and methods

### 2.1. Participants

25 subjects (15 female, 10 male, age = 23.8, SD = 2.2) participated in the experiment. All participants were right-handed (Edinburgh Handedness Inventory), had normal or corrected-to-normal vision and reported no history of neurological or psychiatric diseases. Three participants were excluded after data acquisition – two because of heavy head motion and one due to suspected neurological issues. Accordingly, the final sample comprised 22 individuals (13 female, 9 male, age = 23.6, SD = 2.0). The study protocol was approved by the local ethics committee in accordance with the Declaration of Helsinki, and all participants provided written informed consent.

### 2.2. Stimuli and procedure

Participants performed a visual intensity judgment task in the MRI scanner with constant, minimal lighting distributed equally across the scanner room. The stimuli were presented on a screen (refresh rate 60Hz) located behind the scanner which participants saw via an MR-compatible eye-tracking mirror. Additionally, participants were equipped with two button pads, one for each hand, that were placed on the respective legs. The software used for stimulus presentation was PsychoPy (version 1.64; Peirce et al., 2019). Luminance measurements were performed using an i1Display Pro photometer (X-Rite Pantone, Grand Rapids, USA).

The task consisted of judging which one of two grey discs was brighter. Both stimuli were in the color of the grey of the monitor (R = G = B). The first stimulus was always presented at a luminance of 29.5 cd/m^2^, whereas the second could either have a luminance of 26.6, 28.0, 29.5, 31.0, or 32.7 cd/m^2^. Brightness is a non-linear function of luminance, that is, equal increments in luminance do not correspond to equal increments in brightness (Stevens, 1957, 1966; Poynton, 1993). We chose precisely the aforementioned luminance values since they could be expected to be approximately perceptually equidistant according to the CIE 1976 L* function, given that the white of monitor (R = G = B = max; 68.8 cd/m^2^) is used as reference white for L*. All stimuli were presented against a background which also had the color of the monitor grey and always had a luminance of 16.1 cd/m^2^. We added a white frame (68.8 cd/m^2^) to this background to control for any unsought anchoring effects (Gilchrist and Bonato, 1995; Gilchrist et al., 1999). To diminish possible crispening effects (i.e., facilitated brightness discrimination for stimuli similar in intensity to the background; Takasaki, 1966), a black outline (2 pixel wide) was added to the stimulus (Whittle, 1992).

Trials started with a black fixation dot (0.5° visual angle), followed by an actively elicited or automatically triggered cue (enlarged dot, 0.9° visual angle, 300 ms) which indicated to the participant that the first stimulus would be launched shortly. The strength of the effect of sensory attenuation decreases within a few hundred milliseconds after action execution (Bays et al., 2005; Aliu et al., 2009), thus, we expected sensory attenuation to affect the first of the two stimuli more strongly than the second, as it was closer in time to the button press. Consequently, all imaging and pupillometry analyses will reference to the first stimulus, which was physically identical throughout the experiment and will be referred to as “stimulus of interest” from here on. Subsequently, the stimulus of interest and the comparison stimulus were presented (both 1000 ms and 2° visual angle), separated by a variable inter-stimulus interval (ISI; 1000, 1350, 1650, 2000 ms). Upon offset of the comparison stimulus, participants were prompted with the question “Which stimulus was brighter?” and answered by button press within a time window of 2500 ms. The response was followed by an inter-trial interval (ITI) which defaulted to a duration of 500 ms, with added jitter based on the remaining time from the active button press phase and the response phase (see Fig. 1).

**Figure 1.**
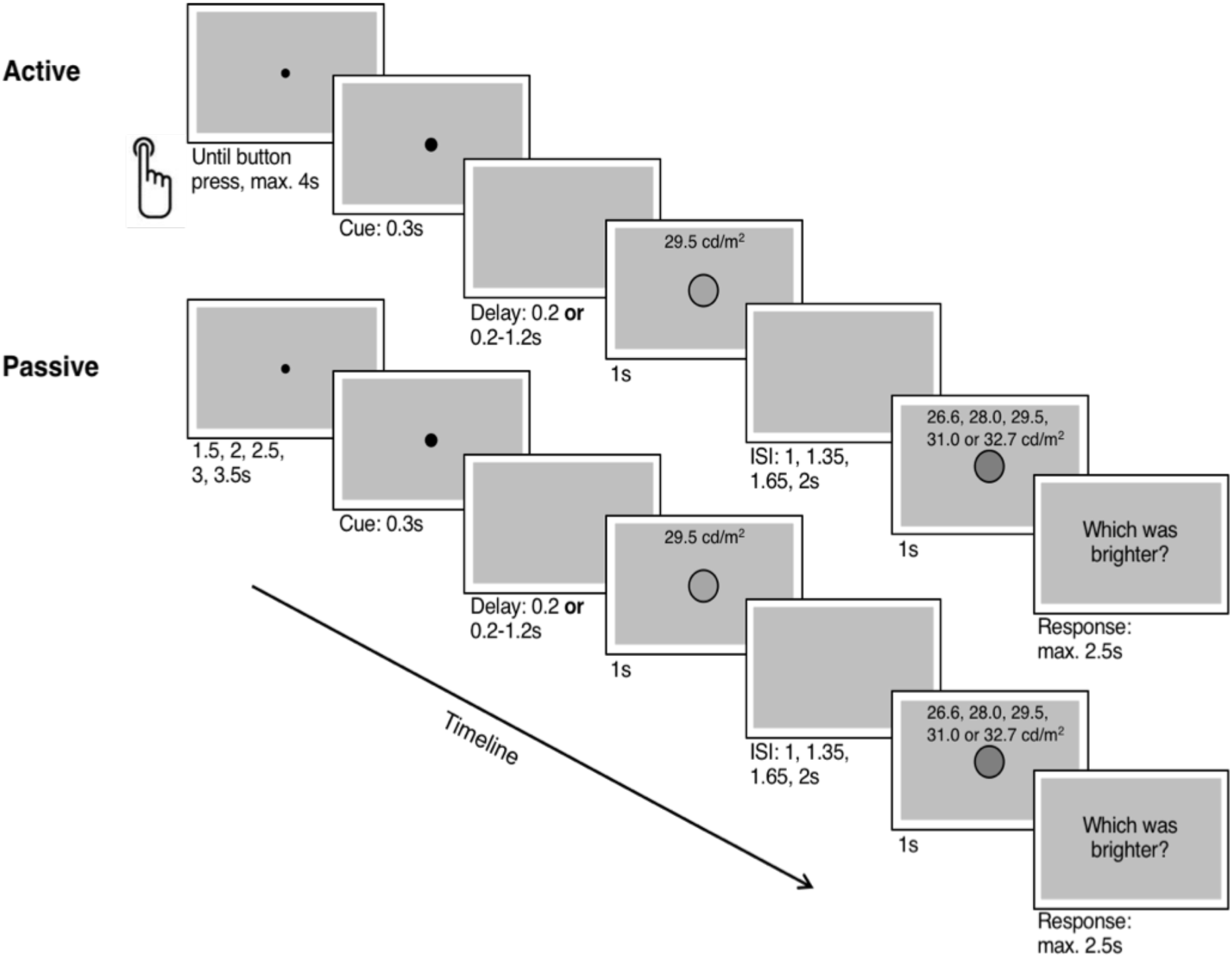
Example of active and passive trials. In active trials (top row), participants performed a button press at any time within 4 seconds after trial onset (i.e., appearance of fixation dot). The button press triggered a cue which was followed by the presentation of the visual stimulus. Stimuli were presented either after a variable delay (unpredictable, 200, 450, 700, 950, 1200 ms) or after a constant period (predictable, 200 ms). Following the stimulus of interest, after a random ISI (1.2-2 seconds) a comparison stimulus was presented. Participants’ task was to report which stimulus they perceived as brighter. In passive trials (bottom row), the trial structure was identical except that participants did not perform button presses, instead cue and stimuli appeared automatically.

The experiment comprised four conditions each of which differed slightly in the details of the trial structure. Trials were manipulated with regard to (1) voluntary generation and (2) the temporal predictability of the stimulus of interest. For the main effect of the efference copy, half of the trials involved a motor action (active) which elicited the stimuli, while no movement was performed in the other half (passive) and stimuli were presented automatically. In active trials, the appearance of the fixation dot indicated the start of a four second time window in which a voluntary button press (right index finger) had to be performed. Participants were instructed to withhold their action for a few milliseconds so that the button press could be consciously prepared and occur in a willed manner, rather than being an automatic mechanism in response to the fixation dot (Rohde and Ernst, 2013). As a direct consequence of the button press, the fixation dot would enlarge and serve as cue signaling the subsequent presentation of the stimuli. In passive trials, no button press was executed. The cue forecasting the upcoming stimulus was identical to the active conditions (enlarged dot) but was generated automatically by the computer. The cue could appear at variable times after trial onset (1500, 2000, 2500, 3000, 3500 ms) to mimic the temporal differences that might occur in the active condition. Notably, the cue indicated a button press (active: the actual button press; passive: a simulated button press performed by the computer) in all conditions and always informed the participant that the action consequence, i.e., the stimulus of interest, would occur after a predictable or unpredictable amount of time in the predictable and unpredictable conditions, respectively. Furthermore, predictability of stimulus onset was manipulated by altering the duration of the time interval between cue offset and onset of the stimulus of interest. In temporally predictable trials, the stimulus of interest was always presented 200 ms after cue offset (thus 500 ms after cue onset). The brief delay was introduced so that participants had the chance to also build up a temporal expectation about stimulus onset in passive trials. In contrast, in unpredictable trials, the onset of the stimulus of interest occurred after a randomly chosen interval (either 200, 450, 700, 950, or 1200 ms, picked with equal probability), relative to cue offset.

Thus, the four conditions were: active predictable (AP), active unpredictable (AU), passive predictable (PP), and passive unpredictable (PU). Each condition (AP, AU, PP, PU) included 60 trials that were evenly distributed across three experimental runs, yielding 240 trials in total (note, that for one participant only 232 trials were collected as data collection was aborted 8 trials prior to the end). Within a run, all conditions were presented and trials were grouped by condition into mini-blocks of 10 trials (2 per run). We pseudo-randomized the order of conditions within a run - so that active and passive conditions alternated - as well as the ISIs and the delay between cue and the stimulus of interest (the latter just for the passive conditions). Prior to each mini-block, participants were visually instructed about the following condition. To assure an identical number of scanned volumes per trial, the differences in trial duration were compensated for in the ITI. The default ITI (500 ms) was extended by the difference between maximum jitter time and presented jitter time in the current trial for (1) all predefined jittered elements (passive button press, unpredictable stimulus appearance, ISI) and (2) all varying elements controlled by the participant (reaction time for active button press and judgment). Consequently, the length of each trial amounted to a total of 12.5 s.

### 2.3. Data acquisition

#### 2.3.1. Functional MRI

MRI Data collection was conducted using a 3 Tesla MR scanner (Siemens Magnetom TIM Trio, Erlangen, Germany) at the Department of Psychiatry and Psychotherapy, Philipps-University Marburg, using a 12-channel head-coil. Participant’s heads were stabilized in order to reduce head motion artifacts. Time courses of functional activation were obtained using a T2*-weighted gradient-echo planar imaging sequence (EPI) sensitive for the blood oxygenation level dependent (BOLD) contrast. Settings were adjusted as follows: echo-planar images, 64 x 64 matrix; 34 slices descending; field of view [FoV] = 192 mm; repetition time [TR] = 1650 ms; echo time [TE] = 25 ms; flip angle = 70°; slice thickness = 4.0 mm, gap size = 15%, and voxel resolution = 3 x 3 x 4.6 mm. Slices were acquired parallel to the intercommissural line (anterior commissure–posterior commissure). During each run of the experimental paradigm, 626 transversal functional whole brain images (including cerebellum) were recorded in descending order. Additionally, a higher resolution T1-weighted volume covering the whole brain was obtained using a magnetization-prepared rapid gradient-echo sequence in sagittal plane (176 slices, TR = 1900 ms, TE = 2.26 ms, FoV = 256 x 256 mm^2^, flip angle 9°, matrix size = 256 x 256 voxels, voxel size = 1.0 x 1.0 x 1.0 mm³).

#### 2.3.2. Eye Tracking

Simultaneously to MR acquisition, we recorded the eye movements and pupil diameter of our participants’ the right eye. Data was collected at a sampling rate of 250 Hz, using an MR-compatible EyeLink 1000 eye-tracker system (SR Research, Osgoode, ON, Canada) that was placed behind the MR scanner, such that participants’ eye movements were recorded via the mirror mounted at the head coil. Prior to each of the three experimental runs, the eye gaze position on the monitor was calibrated using an automated nine-point calibration procedure. The calibration was accepted when the mean error was less than 0.75° of visual angle according to the corresponding validation procedure.

### 2.4. Data analysis

#### 2.4.1. Behavioral data

To analyze participants’ performance in the intensity judgment task, we calculated the proportion of trials in which the stimulus of interest was perceived as darker. Crucially, the stimulus of interest remained physically constant throughout the experiment; luminance values were manipulated only for the comparison stimulus. Thus, if brightness perception was skewed into one direction, this effect would be purely due to perceptual differences. Trials in which participants (1) failed to perform the active button press (in active trials) and/or (2) failed to report their judgment were excluded (1.95 % of trials). Subsequently, the responses of “second stimulus brighter” were calculated for each participant, condition and luminance level of the second (comparison) stimulus. Logistic psychometric functions were fitted using Psignifit 4 (Schütt et al., 2016) for MATLAB (R2014a Mathworks, Sherborn, Massachusetts), implementing a maximum-likelihood estimation. Based on the function, the thresholds and the slopes were derived for each participant and each condition. Thresholds reflect the intensity value at which participants perceive the comparison stimulus as brighter in 50% of the trials, whereas the slopes refer to the participants’ ability to discriminate between the stimuli of different intensities. Here, the measure of main interest were the perceptual thresholds, because they reflect differences in the perceived brightness of the stimuli as a function of the four experimental conditions (Weiss et al., 2011). When comparing the thresholds for conditions (i.e., active and passive), a shift towards the lower intensities (to the left) indicates that, on average, the comparison stimulus was judged as brighter more often than the stimulus of interest, which always had the same luminance. Thus, a leftward shift is associated with sensory attenuation of the first visual event (stimulus of interest), since the second (comparison) stimulus was perceived as brighter in comparison. To statistically analyze the perceptual differences between the four conditions for the estimated thresholds and slopes, 2-by-2 repeated-measures analyses of variance (rmANOVA) were conducted separately for both measures (IBM SPSS Statistics 21). Within-subject factors were suppression (active vs. passive) and predictability (predictable vs. unpredictable) and the significance level was set at p < .05.

#### 2.4.2. Eye tracking

Eye tracking data were used to assure stimuli were properly fixated and to correlate pupil dilation and brain activation. For fixation control, we analyzed the gaze coordinates during the stimulus of interest. A region of interest in which participants had to fixate was defined by a circle around the center of the screen with a radius of 1.75°, thus covering the whole stimulus area while also taking into account the accuracy of the eye-tracker. After performing a drift-correction for each trial (using the gaze position recorded during the 0.3 s central fixation cue), the percentage of samples in which participants’ gaze were inside of our fixation region of interest (ROI) was determined, where missing data (e.g., due to blinks) were classified as being outside the ROI. An rmANOVA with the factors action (active vs. passive) and predictability (predictive vs. unpredictable) was conducted to test for differences in gaze behavior. For the pupil size analysis, we first normalised the pupil size data. To this end, we used all the pupil traces from 500 ms prior to the onset of the stimulus of interest to its offset. We then computed mean and standard deviation across all of these data within each participant (i.e., 1 value per participant for each measure) and subtracted this mean from all data and divided the result by this standard deviation (i.e., normalized pupil data to *z*-scores). Note that thanks to the within-subject design, this normalization does not affect any statistics on the pupillometry data. As main trial-specific measure we used the mean of this normalized pupil data during the presentation of the stimulus of interest (1000 ms) for each trial. These trial-specific estimates of pupil size were used as a covariate in subsequent fMRI analyses to investigate the relation between pupil size and the hemodynamic response. Blinking has been shown to co-occur with button presses (van Dam and van Ee, 2005) and thus systematic differences between active and passive conditions may impact pupil size estimates. We tested for differences in blinking behavior between conditions and found no main effect of action (F(1,21) = 2.230, *p* = .150, *η^2^_p_* = 0.096) and no action*predictability interaction (F(1,21) = 0.001, *p* = .988, *η^2^_p_* < 0.001). There was a main effect of predictability (F(1,21) = 7.853, *p* = .011, *η^2^_p_* = 0.272).

#### 2.4.3. fMRI data

##### Preprocessing

The analysis of MRI data was performed using Statistical Parametric Mapping (SPM12, https://www.fil.ion.ucl.ac.uk/) in Matlab (R2014a Mathworks, Sherborn, Massachusetts). EPI images were realigned to the mean image to correct for head movements. Each individual’s structural scan (T1 weighted) was co-registered to the functional data, segmented and normalized to the standardized Montreal Neurological Institute (MNI) template ICBM152. Using the standardized structural, all EPI images were warped to MNI space (resampled to a voxel size 2×2×2 mm) and smoothed with a full-width at half maximum kernel (8×8×8 mm).

##### 1^st^ level analyses

On the single subject level, a general linear model (GLM) was designed. The design matrix contained 11 regressors: one regressor for each first (stimulus of interest) and second (comparison) stimulus of all experimental conditions (AP_1, AU_1, PP_1, PU_1, AP_2, AU_2, PP_2, PU_2), modeling the duration of stimulus presentation (1 s). Furthermore, the cue, separated for active and passive trials, and participants’ judgment, indicated by a button press, were each included as a single stick function regressor. In addition, the 6 realignment parameters entered the GLM as regressors of no-interest to control for motion-induced artifacts and a high-pass filter was set to a cut-off period of 128s to remove slow frequencies. BOLD responses were modelled by convolving all regressors of interest with the canonical hemodynamic response function (HRF). Based on the parameter estimates, T-contrasts of the first four stimulus regressors (AP_1, AU_1, PP_1, PU_1) against implicit baseline were fed into a flexible-factorial design for the group-level analysis with the factors action (active vs. passive) and predictability (predictable vs. unpredictable). Thus, the group-level analysis solely focusses on the timepoint around the stimulus of interest.

##### 2^nd^ level analysis

As our main interest was sensory attenuation in the visual system, the occipital cortex was selected as region of interest (ROI) using a mask of the occipital lobe in the Wake Forest University (WFU) Pickatlas (Maldjian et al., 2003) based on the Automated Anatomical Labelling (AAL) atlas (Tzourio-Mazoyer et al., 2002). Subsequently to the ROI analysis, a whole brain analysis was conducted to investigate which areas outside of visual cortex might have been involved in attenuating mechanisms in the context of predictability and voluntary action.

##### Contrasts of interest

To investigate the main effects of action and predictability, we contrasted active and passive conditions [(PP + PU) - (AP + AU)] and predictable and unpredictable conditions [(AU + PU) - (AP + PP)], respectively. Finally, we examined the suppression*predictability interaction ([(AU - AP) - (PU - PP)], [(AP - AU) - (PP - PU)]). For all results reported, the significance level was set at p < .001 (uncorrected), corrected for errors of multiple comparisons at the cluster level. We only report clusters below the corrected threshold (*pFWEc* < .05), unless specified otherwise.

##### Correlation with pupil data

For a second, exploratory analysis, a new GLM was set up to account for changes in participant’s pupil size and its association with neural processing. In addition to the regressors from the first GLM, participant’s trial-specific pupil size during the presentation of the stimulus of interest was included as a parametric regressor. Thus, each first stimulus regressor (AP_1, AU_1, PP_1, PU_1) was weighted parametrically by the individual pupil size. T-contrasts of the parametric modulators against implicit baseline were fed into a 2^nd^ level group analysis (flexible factorial design). Here, we tested both positive and negative correlations between pupil size and BOLD response. For this analysis, we employed family wise error correction (FWE) at a significance level of *p* < .05.

## 3. Results

### 3.1. Behavioral results

To determine the effect of action and predictability on brightness perception, we determined the luminance at which the second stimulus was judged equally bright as the stimulus of interest (point of subjective equality – PSE). To this end, we fitted psychometric functions to the fraction of judgements “second stimulus brighter” as function of log luminance of the second stimulus per individual (Fig. 2A). There was a main effect of action (F(1,21) = 4.54, *p* = .045, *η^2^_p_* = 0.178) on the PSE, while we observed no main effect of predictability (F(1,21) = 1.56, *p* = .226, *η^2^_p_* = 0.069) nor an action*predictability interaction (F(1,21) = 0.266, *p* = .611, *η^2^_p_* = 0.013). PSEs were lower in active (*M*: 29.35 cd/m^2^, *SD*: 1.02 cd/m^2^) than in passive (*M*: 29.46 cd/m^2^, *SD*: 1.03 cd/m^2^; Fig. 2B)^1^. Hence, despite substantial between-subject variance, the within-design reveals a subtle but significant tendency to perceive the stimulus of interest as darker in the active than in the passive condition. The ability to distinguish luminance levels between the two stimuli is reflected in the slope of the psychometric functions. We find no main effect of either factor (action: F(1,21) = 0.865, *p* = .363; predictability: F(1,21) = 0.002, *p* = .965); although there was an interaction (F(1,21) = 4.48, *p* = 047; figure 2C), follow-up tests did not show a significant effect for action, neither for predictable (t(21) = 1.32, *p* = .202) nor for unpredictable stimuli (t(21) = 1.33, *p* = .197).

**Figure 2.**
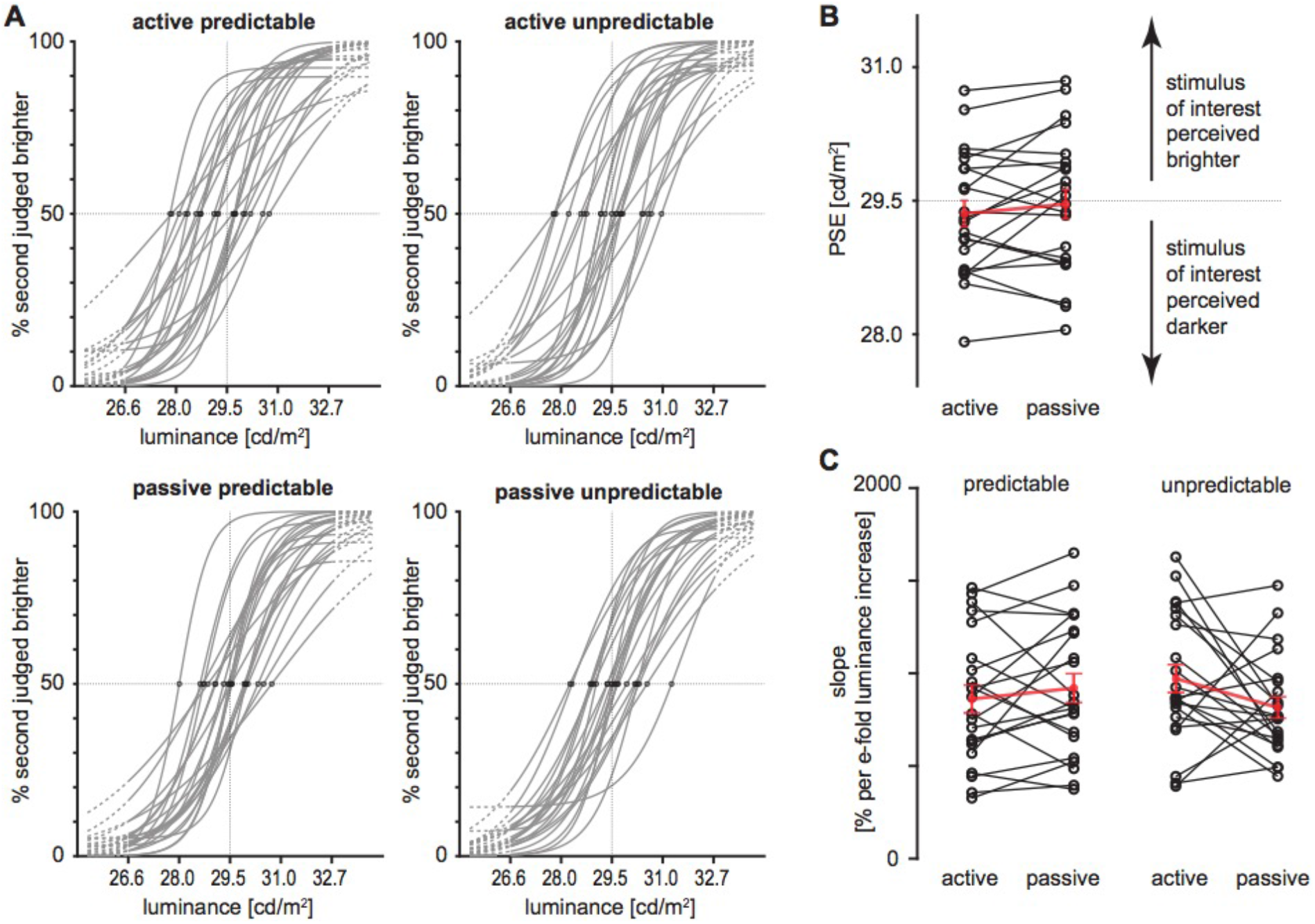
Behavioral data. **A)** Fitted psychometric function per individual (N=22) for the four conditions. Points of subjective equality marked with dots, extrapolation with dotted lines; x-axis logarithmically scaled. **B**) Point of subjective equality averaged over predictabilities, *black lines:* individual participants, *red lines:* mean and SEM. y-axis is logarithmically spaced, dotted line denotes veridical PSE. **C**) Slope of psychometric function for the four conditions, % ‘second judged brighter’ by e-fold increase of luminance; black and red lines as in panel C.

### 3.2. fMRI results

#### ROI analysis

For passive as compared to active conditions (passive > active), we observed bilateral activation in visual cortex, more specifically in calcarine and lingual gyri (x, y, z = 10, −86, 15; T = 4.82, kE = 1004; see Fig. 3A, Table 1) indicating BOLD suppression for active conditions.

**Figure 3.**
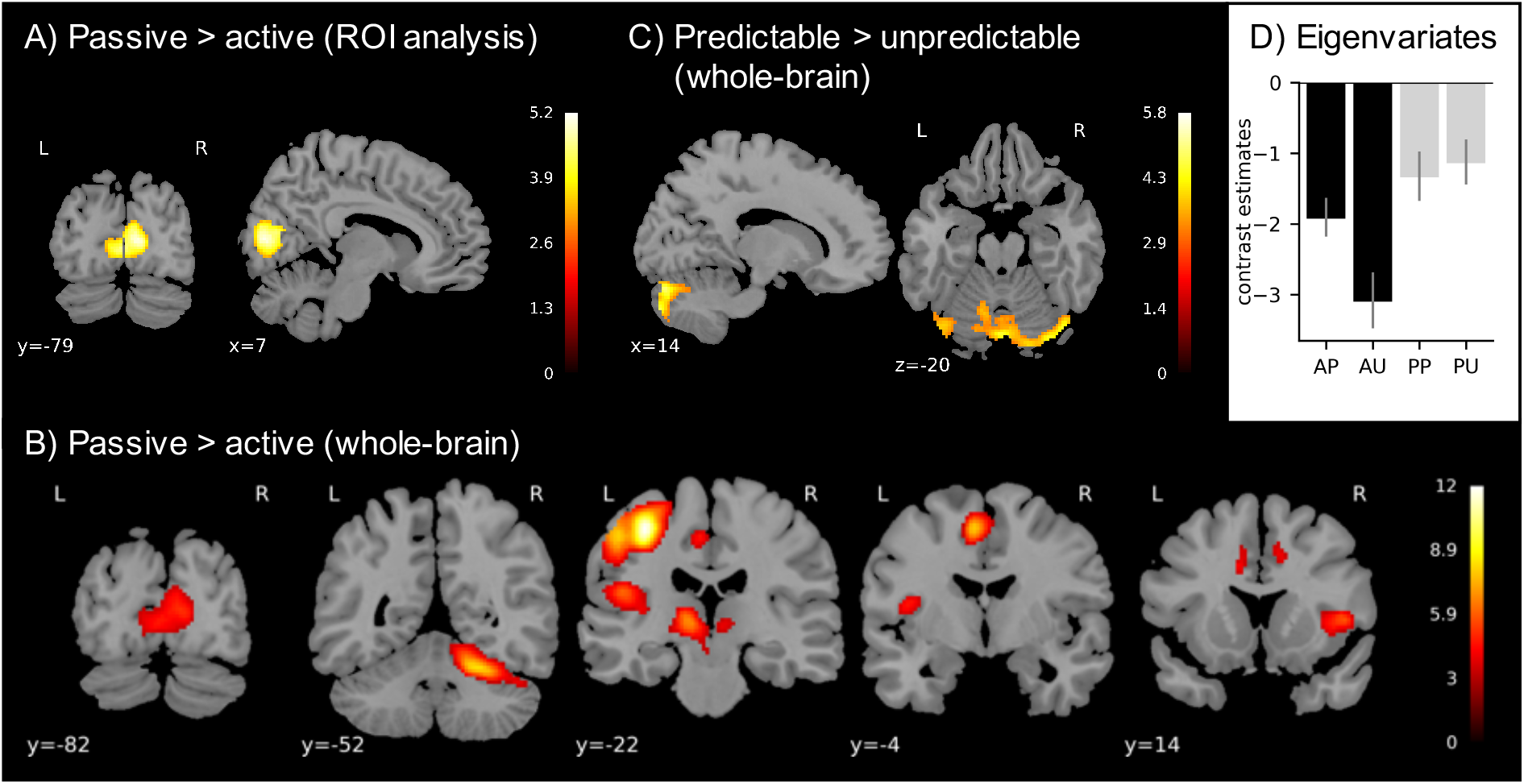
Region of interest and whole-brain analyses. **A.** fMRI results showing BOLD suppression during active conditions (passive > active) in a cluster in visual cortex as assessed by a ROI analysis. **B.** fMRI results for the BOLD suppression in a network of clusters including visual cortex, somatosensory cortex and the cerebellum shown in the whole-brain analysis. **C.** As no main effect of predictability (unpredictable > predictable) was observed, the reverse contrast (predictable > unpredictable) was explored and revealed a cluster in the cerebellum. **D.** Eigenvariates, i.e., the first principal component of the time series, of the cluster with peak activity in Calcarine gyrus extracted from the ROI analysis cluster. Error bars show the standard error of the mean (SEM).

**Table 1.** BOLD suppression for action consequences as compared to identical but externally produced stimuli [(PP + PU) − (AP + AU)]. Coordinates are listed in MNI space. Significance threshold: p < 0.001 uncorrected, pFWEc = 0.05. V1, primary visual cortex; S1, primary somatosensory cortex; S2, secondary somatosensory cortex; IFG, inferior frontal gyrus; PMF, Posterior medial frontal; R, right; L, left.

| Area | Cluster extent | Side | x | y | z | T | k<sub>E</sub> | P<sub>FWEc</sub> |
| --- | --- | --- | --- | --- | --- | --- | --- | --- |
| ROI Analysis |  |  |  |  |  |  |  |  |
| V1 | Calcarine Gyrus | R | 10 | -82 | 10 | 5.19 | 1004 | <0.001 |
|  | Lingual Gyrus | L | -2 | -82 | 4 | 4.67 |  |  |
|  | Calcarine Gyrus | R | 8 | -74 | 14 | 4.49 |  |  |
| Whole-brain Analysis |  |  |  |  |  |  |  |  |
| S1 | Postcentral Gyrus | L | -36 | -22 | 54 | 11.86 | 2249 | <0.001 |
|  | Precentral Gyrus | L | -50 | -24 | 48 | 8.41 |  |  |
|  | Precentral Gyrus | L | -20 | -18 | 68 | 4.23 |  |  |
| Cerebellum | Cerebellum VI | R | 22 | -52 | -22 | 8.3 | 1147 | <0.001 |
|  | Cerebellum VI | R | 20 | -66 | -20 | 4.67 |  |  |
|  | Cerebellum Crus 1 | R | 46 | -50 | -32 | 4.15 |  |  |
| Thalamus | Thalamus | L | -12 | -18 | 2 | 7.96 | 615 | <0.001 |
|  | Thalamus | R | 8 | -16 | 2 | 4.42 |  |  |
|  | Thalamus | R | 2 | -26 | -2 | 3.98 |  |  |
| IFG | Pars Opercularis | R | 46 | 14 | 4 | 5.79 | 520 | <0.001 |
|  | Insula | R | 32 | 24 | 2 | 5.6 |  |  |
|  | Pars Orbitalis | R | 30 | 24 | -12 | 4.94 |  |  |
| S2 | Rolandic operculum | L | -48 | -24 | 20 | 6.3 | 1028 | <0.001 |
|  | Precentral Gyrus | L | -56 | 4 | 28 | 6.18 |  |  |
|  | Precentral Gyrus | L | -38 | -16 | 8 | 5.34 |  |  |
| **PMF** | **SMA** | **L** | **-4** | **-12** | **52** | **8.95** | **1186** | **<0.001** |
|  | MCC | L | -8 | -24 | 48 | 4.34 |  |  |
|  | SMA | L | 8 | 6 | 50 | 4.31 |  |  |
| **V1** | **Calcarine Gyrus** | **R** | **10** | **-82** | **10** | **5.19** | **1179** | **<0.001** |
|  | Lingual Gyrus | L | -2 | -82 | 10 | 4.67 |  |  |
|  | Calcarine Gyrus | R | 8 | -74 | 14 | 4.49 |  |  |

To investigate the influence of predictability on sensory attenuation, conditions of predictable events were subtracted from unpredictable ones (unpredictable > predictable). No clusters of activation were found in visual cortex for this contrast. Similarly, for the action*predictability interaction, reflecting modulatory influences of action generation and temporal predictability on brightness perception, no significant clusters of activation were observed.

As predictability was expected to have an effect, we performed an additional post-hoc analysis to examine whether our manipulation of inserting jittered delays per se worked. To this end, we split our unpredictable trials according to the delay inserted between action and the stimulus of interest (200, 450, 700, 950, 1200 ms) and formed regressors for each delay. Contrasts against implicit baseline were fed into our second level analysis, where we examined the effect of delay. We observed a main effect of delay in the visual cortex (x, y, z = 22, −80, 10; T = 4.04, kE = 164, *p* = .017, uncorrected), with activity linearly increasing as a function of delay (Fig. 4B). This effect was similar in both active and passive conditions; even though activity was overall lower during active trials, both conditions show a linear effect of delay with the lowest activity closest to the action/cue.

**Figure 4.**
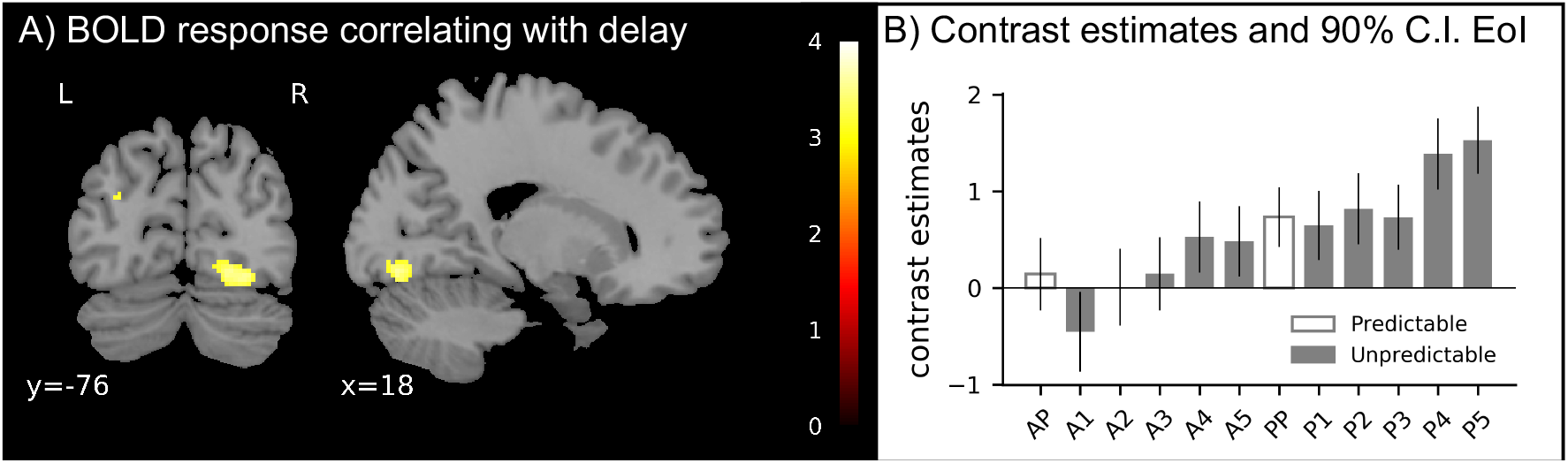
Main effect of delay in a whole-brain analysis. **A.** BOLD activation in a cluster in visual cortex which correlated positively with delay in active and passive conditions. **B.** The bar plot illustrates the mean of extracted eigenvariates (i.e., the first principal component of the ROI’s time series) for active and passive conditions when predictable and unpredictable (separately for all five delay increments) for the cluster peaking in visual cortex [22,-80,-10]. The error bars represent the standard error of the mean (SEM). AP = active predictable, PP = passive predictable, A* = active, P* = passive. The numbers 1 to 5 refer to the delay increments (1 = 1500 ms, 2 = 2000 ms, 3 = 2500 ms, 4 = 3000 ms, 5 = 3500 ms).

#### Whole-brain analysis

In agreement with the ROI analysis, the whole-brain analysis revealed a suppression effect (passive > action) in visual cortex (x, y, z = 10, −82, 10; T = 5.19, kE = 1179; Fig. 3B, Table 1). Additionally, we observed activation in motor-related areas (primary somatosensory: x, y, z = −36, −22, 54, T =11.86, kE = 2249; supplementary motor area (SMA): x, y, z = −4, −12, 52, T =8.95, kE = 1186; cerebellum lobule VI: x, y, z = 22, −52, −22, T = 8.3, kE = 1147) and in the thalamus (x, y, z = −12, −18, 2, T = 7.96, kE = 615).

Also consistent with the ROI analysis, no main effect of predictability (unpredictable > predictable) was observed in the whole-brain analysis, even at a lower threshold of *p* < .001 uncorrected. To further explore the data, the reverse contrast (predictable > unpredictable) was also examined (Fig. 3C, Table 2). For this contrast widely spread activation was observed in the cerebellum involving lobules VIIa (x, y, z = 32, −82, −24, T = 5.78, kE = 3008). Furthermore, this contrast revealed clusters in the superior frontal gyrus (x, y, z = −26, 64, 10, T = 4.66, kE = 210), paracentral gyrus (x, y, z = −8, −24, 78, T = 5.02, kE = 471) and putamen (x, y, z = −16, 12, −12, T = 4.58, kE = 224). The action*predictability interaction revealed one cluster in middle frontal gyrus (x, y, z = 42, 38, 30, T = 5.14, kE = 234), indicating lowest activation for the active unpredictable condition.

**Table 2.**
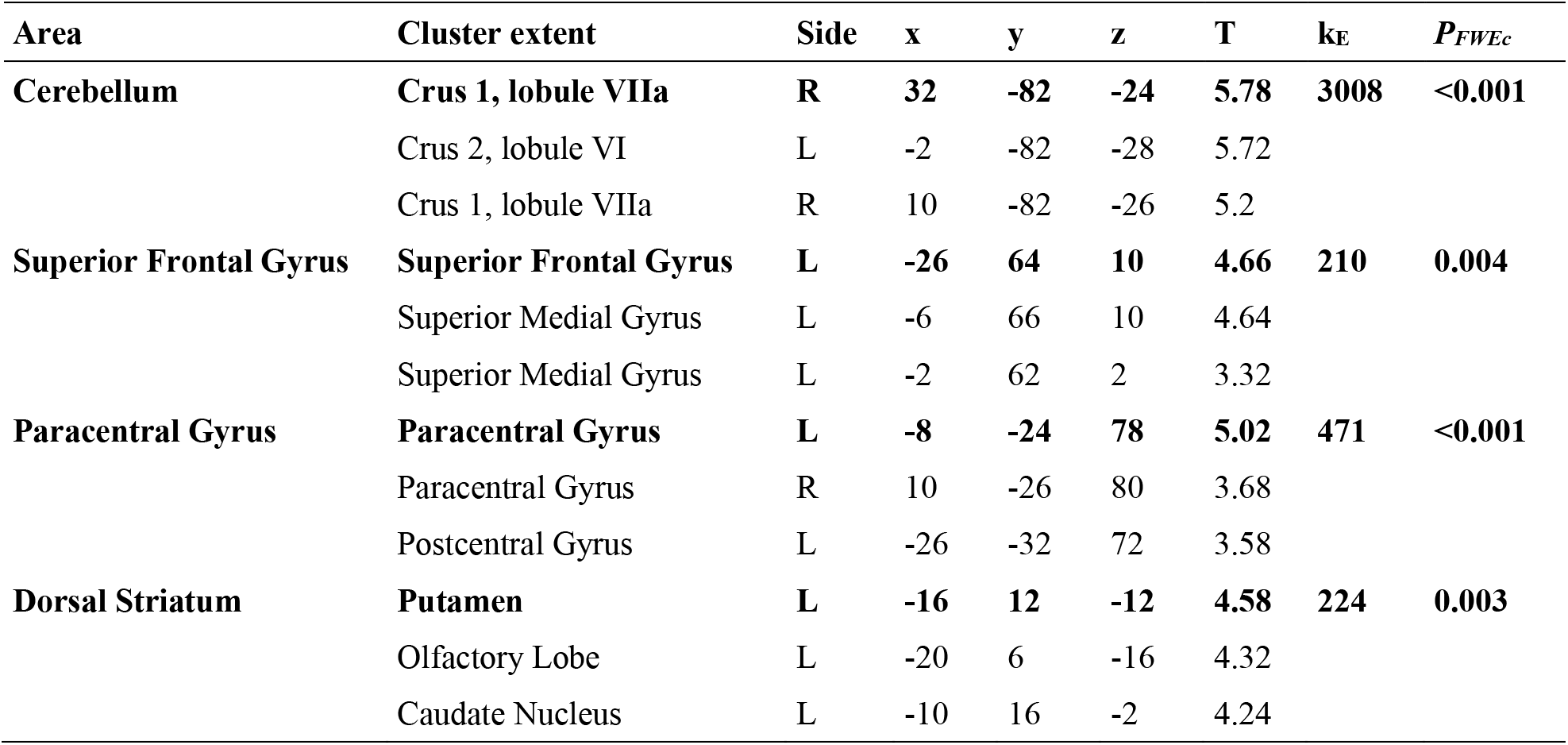
Processing of predictable as compared to unpredictable stimuli [(AP + PP) - (AU - PU)]. Coordinates are listed in MNI space. pFWEc = 0.05. R, right; L, left. Significance threshold: p < 0.004 uncorrected, pFWEc = 0.05. R, right; L, left.

#### Correlation with pupil size

The analysis of pupil size revealed a main effect of action (F(1,21) = 86.01, *p* < .001, *η^2^_p_* = 0.804), such that pupils dilated more strongly in both active (AP: *M* = 0.24; AU: *M* = 0.22) as compared to passive (PP: *M* = −0.20; PU: *M* = −0.35) conditions (Fig. 5A). In addition, we observed a main effect of predictability (F(1,21) = 4.40, *p* = .048, *η^2^_p_* = 0.173) indicating larger pupil size for predictable relative to unpredictable trials. The interaction of action*predictability showed no significant effect but a trend (F(1,21) = 3.91, *p* = .061, *η^2^_p_* = 0.157) thus we tested which condition was driving the difference by comparing pupil diameter between predictable and unpredictable trials in active and passive conditions. We observed that the effect was driven by passive conditions, as in active conditions, diameter did not differ statistically between predictable and unpredictable stimuli (t(21) = 0.24, *p* = .73), whereas pupils were more constricted for passive unpredictable as compared to passive predictable trials (t(21) = 2.80, *p* = .01).

**Figure 5.**
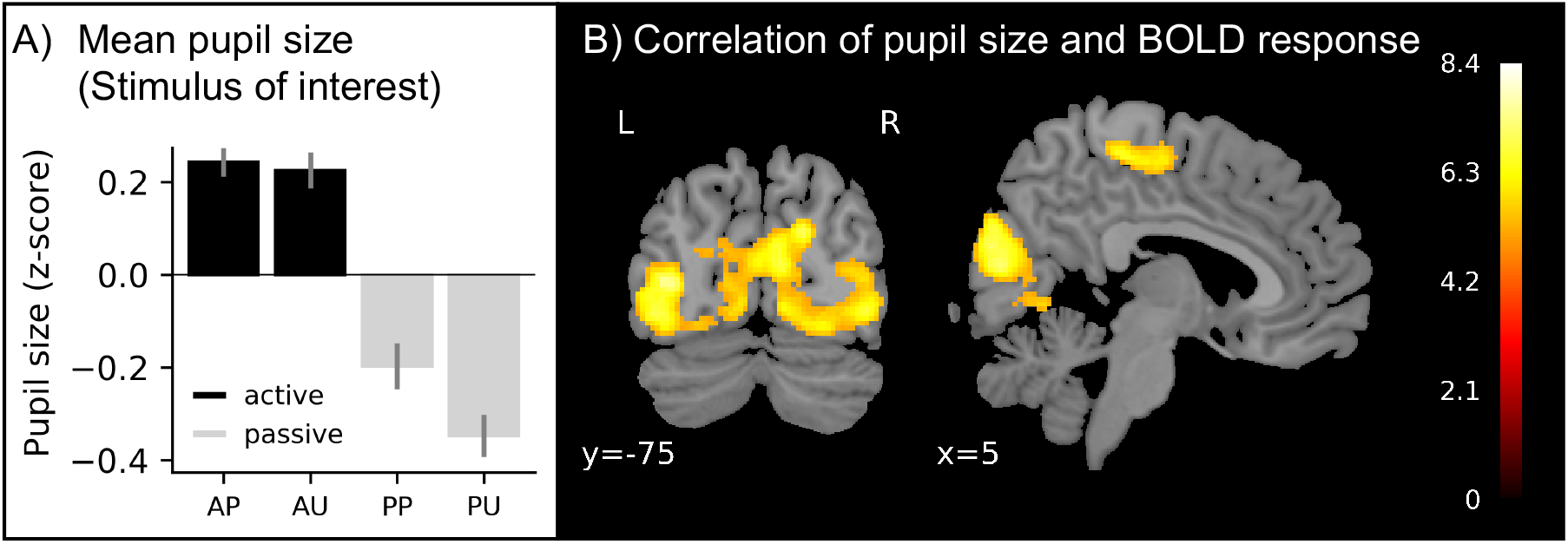
Correlation of pupil size and BOLD response. **A.** Bar plot illustrates z-transformed pupil size averaged across participants separately for each condition. **B.** Activation of brain areas showing a negative linear correlation with pupil size included a network of occipital cortex, pre- and postcentral lobe and rolandic operculum.

In the parametric fMRI analysis with pupil size as a parametric modulator, a negative effect of condition was found: pupil dilation correlated negatively with the hemodynamic response across all conditions. Thus, the smaller the pupil on a given trial, the stronger the BOLD response (Fig. 5B). This association was observed for a widespread cluster in occipital lobe (x, y, z = 3, −84, 15, T = 8.4, kE = 8135), and clusters in pre- and postcentral gyrus (x, y, z = 52, −12, 48, T = 7.5, kE = 3049; x, y, z = −38, −16, 46, T = 8.3, kE = 958) and rolandic operculum (x, y, z = 40, −16, 24, T = 5.7, kE = 43). The results showed no main effect of action, main effect of predictability or action*predictability interaction.

### 3.3. Eye-tracking results

#### Fixation analysis

If the ROI is defined by a radius of 1.75° around the center of the screen, then on average 95.6% of all samples recorded during the presentation of the stimulus of interest indicated that gaze was inside the ROI (second stimulus: 95.1%), where missing data (e.g., due to blinks) were classified as being outside the region of interest. An rmANOVA on the gaze coordinates during the stimulus of interest showed no significant differences between conditions: neither the main effect of action (F(1,21) = 0.130, *p* = 0.72), predictability (F(1,21) = 0.018, *p* = 0.77), nor the action*predictability interaction (F(1,21) = 0.798, *p* = 0.38) reached significance. It is thus very unlikely that our behavioral and fMRI results can be explained by differences in fixation.

## 4. Discussion

Action-based sensory attenuation is a well-known effect in the auditory and tactile modalities, while heterogenous results have been reported in the visual domain. In this fMRI study, we examined whether action-based sensory attenuation can be observed in vision and which roles efference copy and temporal prediction mechanisms play in its generation. The results demonstrate that perceived intensity was lower and neural processing was suppressed in a network including visual, somatosensory and cerebellar brain areas in active as compared to passive conditions. There was no statistically significant effect of predictability, neither for the behavioral nor the neural data. Pupil size was larger in active as compared to passive trials and correlated negatively with BOLD response across all conditions. Overall, these data indicate that sensory attenuation and BOLD suppression are based on action-related rather than temporal predictive mechanisms, possibly related to pupil size.

### 4.1. Suppression of visual cortex activation

Our first main finding is that visual stimuli were perceived as darker and processed with less neural resources when they were actively generated, as compared to identical passively elicited stimuli. Thus, our data suggest that, beyond the auditory and somatosensory systems, sensory attenuation also occurs in the visual modality. This finding is in line with previous work reporting BOLD suppression in visual cortex (Uhlmann et al., inPress, 2020; Straube et al., 2017; Arikan et al., 2019; Pazen et al., 2019; Schmitter et al., 2021) and reduced visual N1 components (Mifsud et al., 2018) for self-generated compared to automatically presented visual stimuli. According to the notion of internal forward models (Wolpert et al., 1995; Miall and Wolpert, 1996), neural processing of action consequences is attenuated because of a cancellation of sensory information that matches the efference copy-based prediction (Blakemore et al., 1998; Shergill et al., 2013). Thus, the suppressed activity we observed in visual cortex for actively as compared to automatically generated stimuli may be a result of such action-based prediction.

An alternative framework that the BOLD suppression may be interpreted in is the “expectation by sharpening” hypothesis (Lee and Mumford, 2003) which is related to the predictive coding theory (for a review see: Summerfield and de Lange, 2014). While this account predicts that predictable stimuli result in reduced aggregate neural activity (Isaacson and Scanziani, 2011), it also assumes a sharper representation of predictable stimulus compared to unpredicted stimuli (Kok et al., 2012). According to the sharpening account, BOLD suppression reflects a deactivation of neurons tuned away from the stimulus by which the activity of neurons coding the actual stimulus is masked. In line with this theory, sharpened sensory representations have also been reported in sensorimotor prediction for stimuli congruent with a hand movement (Yon et al., 2018). In this study, stimuli congruent with a hand movement were decoded with higher accuracy than incongruent action consequences, while at the same time BOLD suppression was observed in voxels tuned away from the stimulus. The analysis presented in the present study cannot adjudicate between the two models. However, while the sharpening theory may explain the BOLD results, it predicts contrasting behavioral effects to what was observed here as the predictive coding framework generally assumes facilitated perception (Mumford, 1992; Friston, 2003).

The behavioral results revealed lower perceptual thresholds in active compared to passive conditions, i.e., the stimulus of interest was perceived as darker when preceded by an active button press. This finding is in line with previous work (Cardoso-Leite et al., 2010; Vasser et al., 2018), parallels the BOLD suppression observed in the fMRI data and is well explained under the cancellation account. Presumably, due to reduced neural processing for stimuli generated by one’s own action the stimulus is represented less faithfully which in turn decreases behavioral performance. Yet the behavioral results presented here stand in contrast to a range of studies reporting no attenuation effect for self-initiated action consequences (Dewey and Carr, 2013; van Kemenade et al., 2016; Yon and Press, 2017; Schwarz et al., 2018) in similar experiments. It is possible that the heterogeneity of stimuli and experimental designs used in the literature, ranging from low level (Schafer and Marcus, 1973; Cardoso-Leite et al., 2010; Gentsch and Schütz-Bosbach, 2011; van Kemenade et al., 2016; Straube et al., 2017; Mifsud et al., 2018) to complex (Hughes and Waszak, 2014) and from discrete to continuously presented (Schmitter et al., 2021) stimuli, may have contributed to the ambiguous findings regarding the attenuation of visual stimuli. Furthermore, studies employed diverse measures such as speed judgments (Dewey and Carr, 2013), contrast discrimination (Roussel et al., 2013), stimulus detection (Cardoso-Leite et al., 2010; Schwarz et al., 2018), delay detections (Uhlmann et al., inPress, 2020; van Kemenade et al., 2016; Straube et al., 2017; Arikan et al., 2019; Pazen et al., 2019) and brightness judgments (Yon and Press, 2017). Further research is necessary to carefully establish the nature of stimuli and behavioral tasks leading to attenuated visual processing.

In line with our hypothesis, we observed an attenuated BOLD response for actively elicited visual stimuli in primary visual cortex. Interestingly, this effect manifested itself in peripheral rather than foveal areas of V1 (Fig 3A and B). At first glance, this finding is counter-intuitive as an effect may be expected in foveal regions representing the stimulus. However, brightness judgments are strongly influenced by the luminance contrast between a stimulus and the background it is presented against, as has been evidenced by effects such as brightness induction (Heinemann, 1955) and anchoring (Gilchrist et al., 1999). Furthermore, luminance contrast has been shown to be represented not only in the lateral geniculate nucleus but also in V1 (Wiesel and Hubel, 1966; Johnson et al., 2001; Kinoshita and Komatsu, 2001; Vinke and Ling, 2020).

In our experiment, luminance contrast was largest at the border between stimulus and background. Thus, we hypothesize that the cluster in primary visual cortex showing BOLD suppression may be a representation of the edge of the stimulus rather than the stimulus as a whole for which a more foveal effect would be expected. In that case, instead of using the luminance to compare the stimulus of interest and the comparison stimulus, participants might rather have based their intensity judgment on the luminance contrast between background and stimulus.

### 4.2. No attenuation of temporally predictable visual stimuli

Unexpectedly, the perceived brightness and neural processing of predictable and unpredictable stimuli did not differ significantly in our experiment. While null findings are no proof for the absence of an effect, they provide the possibility that the effect in question is unlikely to exist. This possibility and its implications will be discussed below.

Our results suggest that the attenuation of behavior and neural processing is not confounded by temporal predictions but may rather be generated by motor-based predictions. This is in line with work from the auditory domain (Klaffehn et al., 2019), which showed attenuated processing of action consequences regardless of whether these were predictable or unpredictable. Combined with the present findings, this may suggest that the prediction of forward models can tolerate some degree of temporal uncertainty and cancellation of matching incoming signals is successful even at delay. However, another recent study evidenced that controlling for temporal predictability can abolish the attenuation of the auditory N1 for self-generated as compared to identical external tones (Kaiser and Schütz-Bosbach, 2018). It is possible that the disparate findings may be related to differences regarding visual stimulation between conditions (constant vs. variable) in the paradigms (Besle et al., 2004; Maddox et al., 2015).

Cortical processing of temporally predictable stimuli is often suppressed (Summerfield et al., 2008; Bendixen et al., 2009; Alink et al., 2010; Todorovic et al., 2011; Kok et al., 2012; John-Saaltink et al., 2015). Here, we manipulated the temporal predictability of visual events by variably delaying the period between cue offset and onset of the stimulus of interest, so that participants were unable to predict stimulus onset in -both actively and passively generated-unpredictable trials. However, participants were aware that a stimulus would certainly be presented on every trial. Thus, absence of significant suppression effects for predictable as compared to unpredictable trials may be explained by the fact that unpredictable conditions were not fully unpredictable in our design. Differences between studies might also be related to the possibility that the role of temporal predictability is not identical across modalities. It has been suggested that temporal predictability of auditory stimuli abolishes the attenuation effect, while visual components did not differ between temporally predictable and unpredictable passive stimuli (Mifsud et al., 2016b). Finally, alongside temporal predictability, further stimulus characteristics (i.e., temporal control, identity prediction) have been proposed to potentially account for the attenuation effects often observed for sensory action outcomes (Hughes et al., 2013). In the present paradigm, stimulus identity did not differ between conditions but, in contrast to active predictable conditions, participants did not have temporal control in passive conditions which may play a role in the results presented here.

### 4.3. Inverse relationship between pupil size and visual cortex activity

According to the pupillary light response effect, decreasing light intensity leads to stronger pupil dilation and increasing light intensity to stronger pupil constriction (Loewenfeld, 1958; Mathôt and Van der Stigchel, 2015). In our experiment, the stimulus of interest remained identical in luminance such that differences in pupil size could not have resulted from its physical properties. Interestingly, we observed that pupil size was larger in the active as compared to the passive conditions. Importantly, recent work suggested that beyond well-known influences of luminance, pupil size is also sensitive to changes in *perceived* luminance (i.e., brightness) (Laeng and Endestad, 2012). Thus, larger pupil sizes in active trials might indicate that participants perceived the stimulus as darker as compared to the identical stimuli in passive trials. This interpretation is in line with our behavioral results which showed that self-generated visual stimuli were perceived as darker and were associated with larger pupil size.

An alternative explanation for the pupil size effect could be the difference in motor action involved in the active and passive conditions as movements, i.e. a button press, can result in pupil size changes, even in the absence of visual stimulation (Richer and Beatty, 1985; Hupe et al., 2009). However, we also observed a negative correlation of pupil size with the BOLD response in visual cortex. Thus, we find that with increasing pupil size, the activation of visual cortex decreases. This association between visual cortex and pupil size supports our hypothesis that the pupil size effect is related to visual processing of the stimulus, rather than to button press-related motion. Note, however, that the correlation was not specific to a certain visual brain area but rather generic for the whole visual cortex. One previous study reported an inverse relationship between pupil size and the response of primary visual cortex (Bombeke et al., 2016) such that (apparently) brighter stimuli were associated with smaller pupil size and a more pronounced C1 component as compared to stimuli perceived as darker. Nevertheless, these results should be interpreted with caution as the measures of pupil size and the C1 component were not correlated and the reported difference in pupil size between conditions was very small (0.02 mm diameter change; Mathôt et al., 2018). As a side note, we find that pupil size is slightly larger when the stimulus is predictable; this is in line with earlier findings that show a larger pupil dilation for less uncertainty about an outcome (Preuschoff et al., 2011).

## 5. Conclusion

In this experiment, behavioral, neuroimaging as well as pupil size results substantiate the existence of sensory attenuation in the visual system. Stimuli elicited by a voluntary button press were perceived as darker, were associated with a suppressed BOLD response in visual cortex and led to smaller pupil size. Interestingly, we demonstrate that pupil size was negatively correlated with the neural response in visual cortex. No significant effect of temporal predictability on the perception or processing of visual stimuli was observed. Our data suggest that sensory attenuation in vision likely relies more on mechanisms based on efference copies than on temporal predictions.

## Data and code availability statement

The data that support the findings of this study will be made openly available in Zenodo.

## Acknowledgements

We wish to thank the Core Unit Brain Imaging for their help with the data collection as well as Jens Sommer for assistance with the implementation of the experimental setup. This work was supported by the Deutsche Forschungsgemeinschaft (DFG, German Research Foundation) – project number 222641018 – SFB/TRR 135 A3 and B4. Benjamin Straube is supported by the German Research Foundation (DFG; STR 1146/15-1, project no. 286893149, and STR 1146/9-2, project no. 429442932).

## Footnotes

1 Note that the psychometric functions were fit to log luminance and arithmetic mean and standard deviation were also determined in log luminance and mapped back to the original values for reporting; that is, in linear units the reported mean is the geometric mean (which is the arithmetic mean in log space).

## References

Alink A, Schwiedrzik CM, Kohler A, Singer W, Muckli L (2010) Stimulus predictability reduces responses in primary visual cortex. Journal of Neuroscience 30:2960–2966.

Aliu SO, Houde JF, Nagarajan SS (2009) Motor-induced suppression of the auditory cortex. Journal of Cognitive Neuroscience 21:791–802.

Arikan BE, van Kemenade BM, Podranski K, Steinsträter O, Straube B, Kircher T (2019) Perceiving your hand moving: BOLD suppression in sensory cortices and the role of the cerebellum in the detection of feedback delays. Journal of Vision 19:4.

Avidan G, Harel M, Hendler T, Ben-Bashat D, Zohary E, Malach R (2002) Contrast Sensitivity in Human Visual Areas and Its Relationship to Object Recognition. Journal of Neurophysiology 87:3102–3116.

Bäß P, Jacobsen T, Schröger E (2008) Suppression of the auditory N1 event-related potential component with unpredictable self-initiated tones: Evidence for internal forward models with dynamic stimulation. International Journal of Psychophysiology 70:137–143.

Bäß P, Widmann A, Roye A, Schröger E, Jacobsen T (2009) Attenuated human auditory middle latency response and evoked 40-Hz response to self-initiated sounds. European Journal of Neuroscience 29:1514–1521.

Bays PM, Wolpert DM, Flanagan JR (2005) Perception of the consequences of self-action is temporally tuned and event driven. Current Biology 15:1125–1128.

Bendixen A, Schroger E, Winkler I (2009) I heard that coming: Event-related potential evidence for stimulus-driven prediction in the auditory system. Journal of Neuroscience 29:8447–8451.

Besle J, Fort A, Delpuech C, Giard M-H (2004) Bimodal speech: early suppressive visual effects in human auditory cortex. Eur J Neurosci 20:2225–2234.

Binda P, Pereverzeva M, Murray SO (2013) Pupil constrictions to photographs of the sun. Journal of Vision 13:8–8.

Blakemore S-J, Frith CD, Wolpert DM (1999a) Spatio-temporal prediction modulates the perception of self-produced stimuli. Journal of Cognitive Neuroscience 11:551–559.

Blakemore S-J, Wolpert DM, Frith CD (1998) Central cancellation of self-produced tickle sensation. Nat Neurosci 1:635–640.

Blakemore S-J, Wolpert DM, Frith CD (1999b) The cerebellum contributes to somatosensory cortical activity during self-produced tactile stimulation. NeuroImage 10:448–459.

Bombeke K, Duthoo W, Mueller SC, Hopf J-M, Boehler CN (2016) Pupil size directly modulates the feedforward response in human primary visual cortex independently of attention. NeuroImage 127:67–73.

Boynton GM, Engel SA, Glover GH, Heeger DJ (1996) Linear Systems Analysis of Functional Magnetic Resonance Imaging in Human V1. J Neurosci 16:4207–4221.

Brown H, Adams RA, Parees I, Edwards M, Friston K (2013) Active inference, sensory attenuation and illusions. Cogn Process 14:411–427.

Cardoso-Leite P, Mamassian P, Schütz-Bosbach S, Waszak F (2010) A new look at sensory attenuation: Action-effect anticipation affects sensitivity, not response bias. Psychol Sci 21:1740–1745.

Cornelissen FW (2006) No Functional Magnetic Resonance Imaging Evidence for Brightness and Color Filling-In In Early Human Visual Cortex. Journal of Neuroscience 26:3634–3641.

Dewey JA, Carr TH (2013) Predictable and self-initiated visual motion is judged to be slower than computer generated motion. Consciousness and Cognition 22:987–995.

Friston K (2003) Learning and inference in the brain. Neural Networks 16:1325–1352.

Frith CD, Blakemore S-J, Wolpert DM (2000) Abnormalities in the awareness and control of action. Phil Trans R Soc Lond B 355:1771–1788.

Gentsch A, Schütz-Bosbach S (2011) I did it: Unconscious expectation of sensory consequences modulates the experience of self-agency and Its functional signature. Journal of Cognitive Neuroscience 23:3817–3828.

Gilchrist A, Kossyfidis C, Bonato F, Agostini T, Cataliotti J, Li X, Spehar B, Annan V, Economou E (1999) An anchoring theory of lightness perception. Psychological Review 106:795–834.

Gilchrist AL, Bonato F (1995) Anchoring of lightness values in center-surround displays. Journal of Experimental Psychology: Human Perception and Performance 21:1427–1440.

Graham N, Sutter A, Venkatesan C (1993) Spatial-frequency- and Orentation-Selectivity of Simple and Complex Channels in Region Segregation. Vision Research 33:1893–1911.

Haggard P, Tsakiris M (2009) The Experience of Agency: Feelings, Judgments, and Responsibility. Curr Dir Psychol Sci 18:242–246.

Haynes J-D, Lotto RB, Rees G (2004) Responses of human visual cortex to uniform surfaces. Proceedings of the National Academy of Sciences 101:4286–4291.

Heinemann EG (1955) Simultaneous brightness induction as a function of inducing- and test-field luminances. Journal of Experimental Psychology 50:89–96.

Hubel DH, Wiesel TN (1968) Receptive fields and functional architecture of monkey striate cortex. The Journal of Physiology 195:215–243.

Hughes G, Desantis A, Waszak F (2013) Mechanisms of intentional binding and sensory attenuation: The role of temporal prediction, temporal control, identity prediction, and motor prediction. Psychological Bulletin 139:133–151.

Hughes G, Waszak F (2014) Predicting faces and houses: Category-specific visual action-effect prediction modulates late stages of sensory processing. Neuropsychologia 61:11–18.

Hupe JM, Lamirel C, Lorenceau J (2009) Pupil dynamics during bistable motion perception. Journal of Vision 9:10–10.

Isaacson JS, Scanziani M (2011) How Inhibition Shapes Cortical Activity. Neuron 72:231–243.

John-Saaltink ES, Utzerath C, Kok P, Lau HC, de Lange FP (2015) Expectation suppression in early visual cortex depends on task set Ben Hamed S, ed. PLoS ONE 10:e0131172.

Johnson EN, Hawken MJ, Shapley R (2001) The spatial transformation of color in the primary visual cortex of the macaque monkey. Nat Neurosci 4:409–416.

Kaiser J, Schütz-Bosbach S (2018) Sensory attenuation of self-produced signals does not rely on self-specific motor predictions. Eur J Neurosci 47:1303–1310.

Kinoshita M, Komatsu H (2001) Neural representation of the luminance and brightness of a uniform surface in the macaque primary visual cortex. Journal of Neurophysiology 86:2559–2570.

Klaffehn AL, Baess P, Kunde W, Pfister R (2019) Sensory attenuation prevails when controlling for temporal predictability of self- and externally generated tones. Neuropsychologia 132:107145.

Kok P, Jehee JFM, de Lange FP (2012) Less is more: Expectation sharpens representations in the primary visual cortex. Neuron 75:265–270.

Laeng B, Endestad T (2012) Bright illusions reduce the eye’s pupil. Proceedings of the National Academy of Sciences 109:2162–2167.

Larsen RS, Waters J (2018) Neuromodulatory correlates of pupil dilation. Front Neural Circuits 12:21.

Lee TS, Mumford D (2003) Hierarchical Bayesian inference in the visual cortex. J Opt Soc Am A 20:1434.

Leube D (2003) The neural correlates of perceiving one’s own movements. NeuroImage 20:2084–2090.

Ling S, Pearson J, Blake R (2009) Dissociation of Neural Mechanisms Underlying Orientation Processing in Humans. Current Biology 19:1458–1462.

Loewenfeld IE (1958) Mechanisms of reflex dilatation of the pupil: Historical review and experimental analysis. Doc Ophthalmol 12:185–448.

Maddox RK, Atilgan H, Bizley JK, Lee AK (2015) Auditory selective attention is enhanced by a task-irrelevant temporally coherent visual stimulus in human listeners. eLife 4:e04995.

Maldjian JA, Laurienti PJ, Kraft RA, Burdette JH (2003) An automated method for neuroanatomic and cytoarchitectonic atlas-based interrogation of fMRI data sets. NeuroImage 19:1233–1239.

Martikainen MH (2004) Suppressed responses to self-triggered sounds in the human auditory cortex. Cerebral Cortex 15:299–302.

Mathôt S, Fabius J, Van Heusden E, Van der Stigchel S (2018) Safe and sensible preprocessing and baseline correction of pupil-size data. Behav Res 50:94–106.

Mathôt S, Van der Stigchel S (2015) New light on the mind’s eye: The pupillary light response as active vision. Curr Dir Psychol Sci 24:374–378.

Miall RC, Wolpert DM (1996) Forward models for physiological motor control. Neural Networks 9:1265–1279.

Mifsud NG, Beesley T, Watson TL, Elijah RB, Sharp TS, Whitford TJ (2018) Attenuation of visual evoked responses to hand and saccade-initiated flashes. Cognition 179:14–22.

Mifsud NG, Beesley T, Watson TL, Whitford TJ (2016a) Attenuation of auditory evoked potentials for hand and eye-initiated sounds. Biological Psychology 120:61–68.

Mifsud NG, Oestreich LKL, Jack BN, Ford JM, Roach BJ, Mathalon DH, Whitford TJ (2016b) Self-initiated actions result in suppressed auditory but amplified visual evoked components in healthy participants: Neural response to self-initiated sensory stimuli. Psychophysiol 53:723–732.

Mumford D (1992) On the computational architecture of the neocortex. Biological Cybernetics 66:241–251.

Naber M, Nakayama K (2013) Pupil responses to high-level image content. Journal of Vision 13:7–7.

Pazen M, Uhlmann L, van Kemenade BM, Steinsträter O, Straube B, Kircher T (2019) Predictive perception of self-generated movements: Commonalities and differences in the neural processing of tool and hand actions. NeuroImage:116309.

Peirce J, Gray JR, Simpson S, MacAskill M, Höchenberger R, Sogo H, Kastman E, Lindeløv JK (2019) PsychoPy2: Experiments in behavior made easy. Behav Res 51:195–203.

Penacchio O, Otazu X, Dempere-Marco L (2013) Neuronal population mechanisms of lightness perception. PLOS ONE 8:1–14.

Poynton CA (1993) “Gamma” and its disguises: The nonlinear mappings of intensity in perception, CRTs, film, and video. SMPTE Journal:10.

Preuschoff K, t’Hart BM, Einhäuser W (2011) Pupil dilation signals surprise: evidence for noradrenaline’s role in decision making. Front Neurosci 5.

Richer F, Beatty J (1985) Pupillary dilations in movement preparation and execution. Psychophysiology 22:204–207.

Rohde M, Ernst MO (2013) To lead and to lag – Forward and backward recalibration of perceived visuo-motor simultaneity. Front Psychology 3.

Roussel C, Hughes G, Waszak F (2013) A preactivation account of sensory attenuation. Neuropsychologia 51:922–929.

Sanmiguel I, Todd J, Schröger E (2013) Sensory suppression effects to self-initiated sounds reflect the attenuation of the unspecific N1 component of the auditory ERP: Auditory N1 suppression: N1 components. Psychophysiol 50:334–343.

Sato A (2008) Action observation modulates auditory perception of the consequence of others’ actions. Consciousness and Cognition 17:1219–1227.

Sato A (2009) Both motor prediction and conceptual congruency between preview and action-effect contribute to explicit judgment of agency. Cognition 110:74–83.

Schafer EWP, Marcus MM (1973) Self-stimulation alters human sensory brain responses. Science 181:175–177.

Schmitter CV, Steinsträter O, Kircher T, van Kemenade BM, Straube B (2021) Commonalities and differences in predictive neural processing of discrete vs continuous action feedback. NeuroImage 229:117745.

Schütt HH, Harmeling S, Macke JH, Wichmann FA (2016) Painfree and accurate Bayesian estimation of psychometric functions for (potentially) overdispersed data. Vision Research 122:105–123.

Schwarz KA, Pfister R, Kluge M, Weller L, Kunde W (2018) Do we see it or not? Sensory attenuation in the visual domain. Journal of Experimental Psychology: General 147:418–430.

Shergill SS, White TP, Joyce DW, Bays PM, Wolpert DM, Frith CD (2013) Modulation of somatosensory processing by action. NeuroImage 70:356–362.

Stevens SS (1957) On the psychophysical law. Psychological Review 64:153–181.

Stevens SS (1966) Duration, luminance, and the brightness exponent. Perception & Psychophysics 1:96–100.

Straube B, van Kemenade BM, Arikan BE, Fiehler K, Leube DT, Harris LR, Kircher T (2017) Predicting the multisensory consequences of one’s own action: BOLD suppression in auditory and visual cortices Ahveninen J, ed. PLoS ONE 12:e0169131.

Summerfield C, de Lange FP (2014) Expectation in perceptual decision making: neural and computational mechanisms. Nat Rev Neurosci 15:745–756.

Summerfield C, Trittschuh EH, Monti JM, Mesulam M-M, Egner T (2008) Neural repetition suppression reflects fulfilled perceptual expectations. Nat Neurosci 11:1004–1006.

Takasaki H (1966) Lightness change of grays induced by change in reflectance of gray background. J Opt Soc Am 56:504.

Todorovic A, van Ede F, Maris E, de Lange FP (2011) Prior expectation mediates neural adaptation to repeated sounds in the auditory cortex: An MEG study. Journal of Neuroscience 31:9118–9123.

Tzourio-Mazoyer N, Landeau B, Papathanassiou D, Crivello F, Etard O, Delcroix N, Mazoyer B, Joliot M (2002) Automated anatomical labeling of activations in SPM using a macroscopic anatomical parcellation of the MNI MRI single-subject brain. NeuroImage 15:273–289.

Uhlmann L, Pazen M, Kemenade BM, Steinsträter O, Harris LR, Kircher T, Straube B (2020) Seeing your own or someone else’s hand moving in accordance with your action: The neural interaction of agency and hand identity. Hum Brain Mapp 41:2474–2489.

Uhlmann L, Pazen M, van Kemenade BM, Kircher T, Straube B (inPress) Neural correlates of self-other distinction in patients with schizophrenia spectrum disorders: The roles of agency and hand identity. Schizophrenia Bulletin.

van Dam LCJ, van Ee R (2005) The role of (micro)saccades and blinks in perceptual bi-stability from slant rivalry. Vision Research 45:2417–2435.

van Kemenade BM, Arikan BE, Kircher T, Straube B (2016) Predicting the sensory consequences of one’s own action: First evidence for multisensory facilitation. Atten Percept Psychophys 78:2515–2526.

Vasser M, Vuillaume L, Cleeremans A, Aru J (2018) Waving goodbye to contrast: Self-generated hand movements attenuate visual sensitivity. Neuroscience.

Vinke LN, Ling S (2020) Luminance potentiates human visuocortical responses. Journal of Neurophysiology 123:473–483.

Weiss C, Herwig A, Schütz-Bosbach S (2011) The self in action effects: Selective attenuation of self-generated sounds. Cognition 121:207–218.

Whittle P (1992) Brightness, discriminability and the “Crispening Effect.” Vision Research 32:1493–1507.

Wiesel TN, Hubel DH (1966) Spatial and chromatic interactions in the lateral geniculate body of the rhesus monkey. Journal of Neurophysiology 29:1115–1156.

Wolpert D, Ghahramani Z, Jordan M (1995) An internal model for sensorimotor integration. Science 269:1880–1882.

Yon D, Gilbert SJ, de Lange FP, Press C (2018) Action sharpens sensory representations of expected outcomes. Nat Commun 9:4288.

Yon D, Press C (2017) Predicted action consequences are perceptually facilitated before cancellation. Journal of Experimental Psychology: Human Perception and Performance 43:1073–1083.

